# Shoot-to-root translocation of the jasmonate precursor 12-oxo-phytodienoic acid (OPDA) coordinates plant growth responses following tissue damage

**DOI:** 10.1101/517193

**Authors:** Adina Schulze, Marlene Zimmer, Stefan Mielke, Hagen Stellmach, Charles W. Melnyk, Bettina Hause, Debora Gasperini

**Affiliations:** Department of Molecular Signal Processing, Leibniz Institute of Plant Biochemistry, 06120 Halle (Saale), Germany; Department of Cell and Metabolic Biology, Leibniz Institute of Plant Biochemistry, 06120 Halle (Saale), Germany; Department of Plant Biology, Swedish University of Agricultural Sciences, 75651 Uppsala, Sweden

**Author notes:** Corresponding author: Debora Gasperini.

**Keywords:** Hormone translocation, phloem, OPDA, jasmonate, root growth, grafting

## Abstract

Multicellular organisms rely upon the movement of signaling molecules across cells, tissues and organs to communicate among distal sites. In plants, herbivorous insects, necrotrophic pathogens and mechanical wounding stimulate the activation of the jasmonate (JA) pathway, which in turn triggers the transcriptional changes necessary to protect plants against those challenges, often at the expense of growth. Although previous evidence indicated that JA species can translocate from damaged into distal sites, the identity of the mobile compound(s), the tissues through which they translocate and the consequences of their relocation remain unknown. Here, we demonstrated that endogenous JA species generated after shoot injury translocate to unharmed roots via the phloem vascular tissue in *Arabidopsis thaliana*. By wounding wild-type shoots of chimeric plants and by quantifying the relocating compounds from their JA-deficient roots, we uncovered that the JA-Ile precursor 12-oxo-phytodienoic acid (OPDA) is a mobile JA species. Our data also showed that OPDA is a primary mobile compound relocating to roots where, upon conversion to the bioactive hormone, it induces JA-mediated gene expression and root growth inhibition. Collectively, our findings reveal the existence of long-distance transport of endogenous OPDA which serves as a communication molecule to coordinate shoot-to-root responses, and highlight the importance of a controlled distribution of JA species among organs during plant stress acclimation.

## INTRODUCTION

The existence of complex multicellular organisms would not be possible without efficient communication among distal tissues and organs. In higher plants, the coordination of developmental and environmental responses relies upon the translocation of signal molecules such as phytohormones from one part of the plant to another [1]. Insect herbivory or mechanical wounding trigger an increase in jasmonic acid (JA) and the active hormone jasmonoyl-L-isoleucine (JA-Ile) not only at the site of damage but also in distal, unharmed tissues [2, 3]. Consequently, JA-Ile perception and signaling lead to the activation of defense responses and inhibition of growth across distal tissues to promote plant fitness [4]. Studies in *Arabidopsis thaliana* (Arabidopsis) have shown that wounded leaves emit at least two kinds of signals to alert distal organs: a rapidly transmitted one that travels from leaf-to-leaf [3, 5-7] and an additional one involving the translocation of JA-Ile or its precursors along a basipetal (downwards) direction [8]. The rapid leaf-to-leaf electrical signal can be measured as wound activated surface potentials (WASPs) and is mediated by several clade 3 GLUTAMATE RECEPTOR-LIKE (GLR) genes that stimulate distal hormone production [5-7]. Concomitantly, micrografting experiments between wild-type (wt) scions and JA-deficient rootstocks have demonstrated the existence of a wound-induced shoot-to-root translocation of endogenous JA species that is independent of electrical signals [8]. However, it is still largely unclear which are the endogenously-produced mobile JA forms, what are their transport routes and what are their effects in distal organs.

JA-Ile and its precursors (hereafter referred to as jasmonates, JAs) derive from oxygenated polyunsaturated fatty acids (oxylipins) predominantly through the plastidial octadecanoic pathway (Supplemental Figure 1) [4, 9]. The enzymatic oxygenation of α-linolenic acid (18:3) or hexadecatrienoic acid (16:3) forms the corresponding hydroperoxides 13(S)-hydroperoxy-octadecatrienoic acid (13-HPOT) and 11(S)-hydroperoxy-hexadecatrienoic acid (11-HPHT). 13-HPOT and 11-HPHT are then rearranged by the ALLENE OXIDE SYNTHASE (AOS) to form allene oxides (13S)-12,13-epoxy-octadecatrienoic acid (12,13-EOT) and (11S)-10,11-epoxy-octadecatrienoic acid (10,11-EOT). Because *AOS* is encoded by a single copy gene in Arabidopsis, the *aos* knock-out mutant lacks all compounds downstream from 13-HPOT and 11-HPHT and is hence deficient in all JA-mediated responses (Supplemental Figure 1) [10]. Due to their epoxide ring, 12,13-EOT and 10,11-EOT are extremely unstable compounds [11], which are rapidly converted into *cis*-12-oxo-phytodienoic acid (OPDA) and dinor-oxo-phytodienoic acid (dnOPDA) [12]. OPDA and dnOPDA then translocate from plastids to peroxisomes through the ABC transporter COMATOSE (CTS) [13], where they are reduced to 3-oxo-2-(2-(Z)-pentenyl)-cyclopentane-1-octanoic (OPC-8) and hexanoic (OPC-6) acids, respectively. OPC-8 and OPC-6 undergo several rounds of β-oxidation to generate 3-oxo-2-(2-(Z)-pentenyl)-cyclopentane-1-butanoic acid (OPC-4) and finally JA. Furthermore, OPDA can be converted to dnOPDA to form 4,5-didehydrojasmonate (4,5-ddh-JA) and thereafter JA, via an OPDA Reductase 3 (OPR3)-independent shunt pathway which contributes to a minor fraction of overall JA production [14]. Upon export to the cytoplasm, JA is conjugated to isoleucine (Ile) [15]. Increases in JA-Ile levels trigger the CORONATINE-INSENSITIVE 1 (COI1)-dependent degradation of JASMONATE ZIM DOMAIN (JAZ) transcriptional repressors. JAZ degradation results in the de-repression of transcription factors that initiate JA-dependent transcriptional reprogramming such as the induction of *JASMONATE ZIM-DOMAIN 10* (*JAZ10*) and ultimately promote plant stress acclimation [16-19].

**Figure 1.**
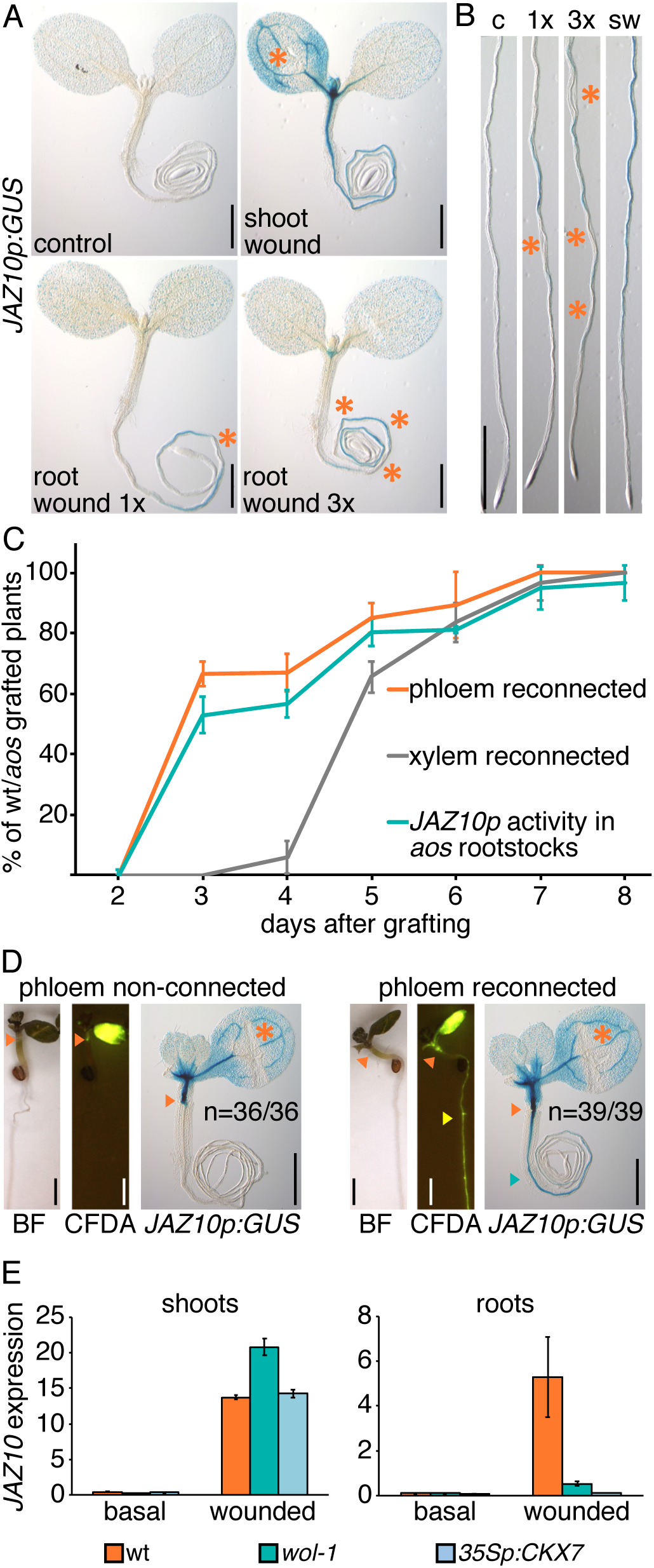
Wound-induced jasmonates translocate from damaged shoots into undamaged roots through the phloem. **(A-B)** Histochemical detection of *JAZ10p:GUS* reporter activity in 5 d old wt seedlings in control (c) and following shoot wounding (sw), or single (1x) and triple (3x) root wounding. Detection of GUS activity was performed 3 h after wounding, damage sites are indicated with orange asterisks. (**B)** Root close-ups of treatments from **A**. Each image is a representative sample of 17-31 biological replicates. Scale bars (**A-B**) = 0.5 mm. **(C)** Time course of phloem and xylem CFDA connectivity assays in wt/*aos* grafts. Following scion wounding, shoot-to-root translocation of jasmonates was tested by *JAZ10p:GUS* reporter induction in the JA-deficient *aos* rootstocks. Phenotype scoring is shown in Supplemental Figure 2. Bars represent means from 3 biological replicates (±SD), each consisting of 9-17 grafted plants. Ungrafted wt was used as control for all time points and assays (n = 10) and exhibited 100% rates. **(D)** Individual wt/*aos* grafts assayed consecutively for phloem connectivity followed by *JAZ10p:GUS* activity after scion wounding at 3 d after grafting. Orange arrowheads indicate graft junctions, yellow arrowhead indicates CFDA staining in the rootstock, blue arrowhead shows GUS reporter activity in the rootstock, bright field (BF). Scale bars = 1 mm. **(E)** Quantitative RT-PCR (qRT-PCR) of *JAZ10* expression in shoots and roots under basal conditions and 1 h after cotyledon wounding of wt, *wol-1*, and *35Sp:CKX7* 5 d old seedlings. *JAZ10* transcript levels were normalized to those of *UBC21*. Bars represent the means of three biological replicates (±SD), each containing a pool of organs from ∼60 seedlings.

Although JA and JA-Ile are abundant in many cells after wounding [20] and thus possible mobile candidates, conclusive evidence of which are the endogenously-produced jasmonate species translocating across long-distances is still missing. Exogenous treatments of labeled OPC-8, JA, or JA-Ile have implied translocation of all tested compounds from the sites of application to distal tissues in different plant species [21-25]. Grafting experiments in tomato (*Solanum lycopersicum*) and wild tobacco (*Nicotiana attenuata*) between wt and JA-deficient mutants have indicated acropetal (upwards) translocation of wound-induced oxylipins including JA and JA-Ile [26, 27]. Studies in barrel medic (*Medicago truncatula*) further support acropetal translocation of JA species [28], although contrasting evidence has also been reported for tomato [29]. Notably, by using an inducible system in Arabidopsis, no evidence was found of wound-induced leaf-to-leaf translocation of JA and JA-Ile [3], leaving open the possibility of other JA precursors being mobile. Overall, the available evidence suggests that the identity of the long-distance transmitted signal might be specific to the direction considered (acropetal, basipetal or leaf-to-leaf) and may vary across plant species.

Here, we investigated which are the shoot-to-root transport routes of wound produced oxylipins; what are the endogenous mobile compounds; and what are the physiological consequences of oxylipin translocation to roots. By coupling grafting experiments to hormone profiling, we uncovered that following wounding, shoot-produced JA species move basipetally through the phloem and restrain root growth. Moreover, we unraveled that the JA-Ile precursor OPDA is a mobile oxylipin mediating shoot-to-root coordination of JA responses in a COI1-dependent manner. Our findings extend previous knowledge on the transport of jasmonates in vascular plants, which have important implications for the regulation of plant growth during stress responses.

## RESULTS AND DISCUSSION

### Jasmonates translocate from wounded shoots into undamaged roots via the phloem

Mechanical wounding of seedling cotyledons triggers JA-mediated gene transcription at the site of damage as well as in below-ground tissues [30]. This can be visualized by the activation of a JA-responsive *JAZ10p:GUS* reporter, which is expressed at very low levels under control conditions (Fig. 1A) [8]. Reciprocal grafting experiments between the wt and the JA-deficient *aos* mutant uncovered that following shoot wounding, endogenous oxylipins translocate to roots to activate *JAZ10p:GUS* expression in young seedlings [8]. In contrast to shoot wounding, mechanical damage inflicted by either single or triple crushes to roots induced *JAZ10p:GUS* expression near the sites of damage, but failed to induce the reporter in aboveground tissues (Fig. 1A-B). This could be due to a smaller proportion of crushed cells and consequent lower amount of oxylipins produced in root tissues compared to shoots, or to a less efficient acropetal long-distance signal in Arabidopsis. Given the much weaker effect observed for wound-triggered root-to-shoot signaling, we focused on the basipetal shoot-to-root direction.

Due to its basipetal flow, it is likely that the shoot-to-root translocation of oxylipins occurs through the phloem, the vascular component that distributes photoassimilates from source to sink tissues. To test this, we monitored phloem and xylem connectivity re-establishment following micrografting of wt scions onto *aos* rootstocks by carboxyfluorescein diacetate (CFDA) assays [31] and correlated them with *JAZ10p:GUS* root induction after wounding (Fig. 1C; Supplemental Figure 2). Because *aos* is JA deficient, activation of *JAZ10p:GUS* expression in *aos* rootstocks can occur only if JA species downstream of 13-HPOT translocate from wounded wt scions (Supplemental Figure 1) [8, 30]. The dynamics of wt/*aos* vascular re-connections were similar to the ones reported for wt/wt combinations [31]: 67 ± 4 % of plants exhibited phloem reconnection at 3 day (d) post grafting, which proceeded until completion at 7 d after grafting. In contrast, xylem reconnection started later with a significant increase in CFDA transport between 4 and 5 d until full completion at 8 d post grafting. The differential reconnection dynamics between phloem and xylem tissues provided us with the opportunity to assay oxylipin transport when only the phloem is connected (3 d post grafting) or when both tissues are reconnected (> 4 d after grafting). We hence monitored wound induced reporter expression in rootstocks of wt/*aos* plants during graft formation and observed that at 3 d post grafting, when only the phloem but not the xylem was connected, 53 ± 6 % of plants showed reporter root induction (Fig. 1C). The activation of reporter expression in wounded wt/*aos* grafts was not a consequence of translocation of the GUS protein or its mRNA from the scion into the rootstock as wounding of wt scions harboring the *JAZ10p:GUS* reporter onto wt rootstocks lacking the reporter did not produce any staining in the rootstock (Supplemental Figure 3). The correlation coefficient between phloem reconnection and oxylipin translocation measured as root reporter activity during the entire time-course was R = 0.99 (p-value = 1.23e^-5^). We next assayed individual seedling consecutively for both phloem connectivity and root reporter induction at 3 d post grafting, when phloem connectivity occurred in about 60 % of individuals (Fig. 1D). Seedlings which lacked phloem connectivity also failed to induce root reporter expression (n = 36/36), and conversely, individuals with re-established phloem flow exhibited a full penetrance of root reporter induction (n = 39/39) (Fig. 1D). Although our assays could not exclude an additional transport route for shoot-produced oxylipins, they clearly indicated that phloem re-connection after grafting was sufficient to translocate them to roots.

**Figure 2.**
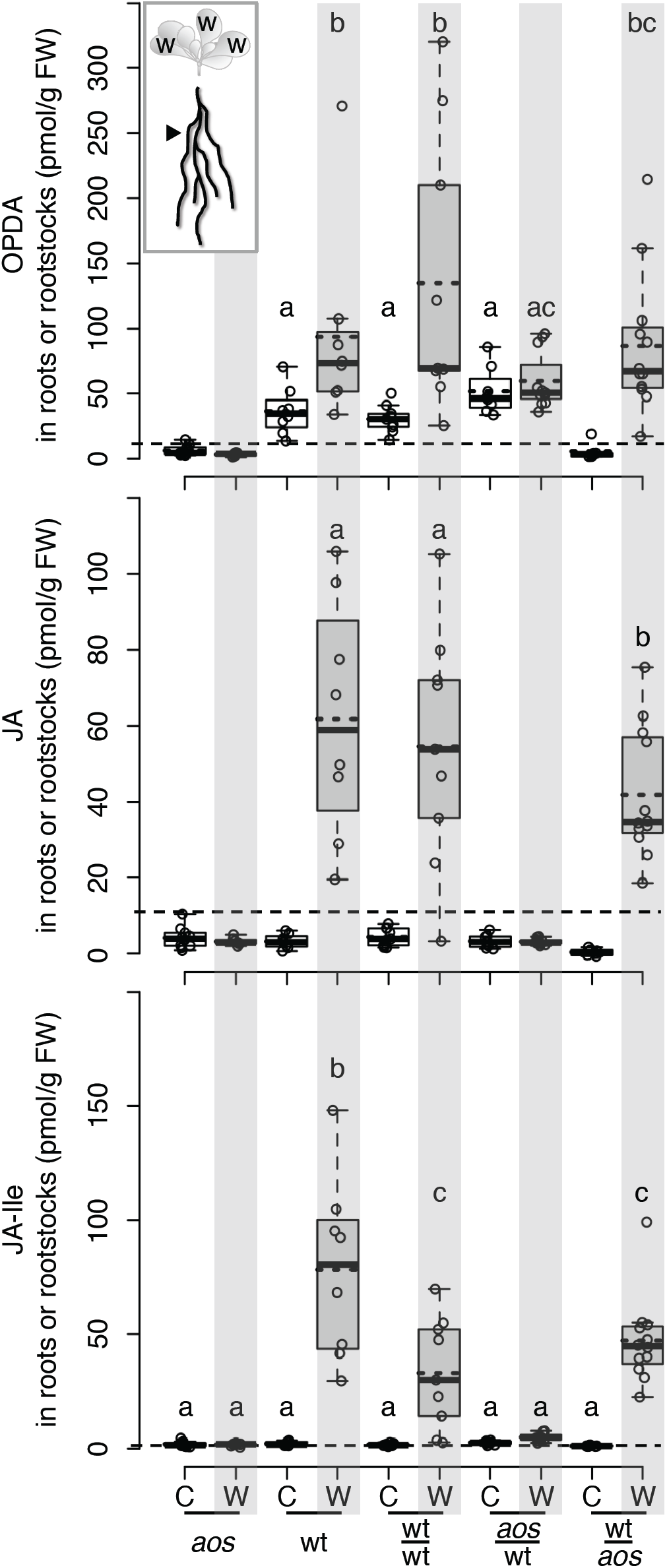
Endogenous OPDA translocates from wounded shoots into unwounded roots. Box plot summary of OPDA, JA, and JA-Ile levels in roots of indicated ungrafted (*aos*, wt) and grafted (wt/wt, *aos*/wt, wt/*aos*) genotypes at control conditions (C) or 1 h following shoot wounding (W). Circles depict biological replicates (7-10 per treatment), each consisting of roots from 15-20 individuals, medians and means are represented inside the boxes by solid black and dotted lines respectively. Values below the limit of quantification (LOQ) indicated by dotted lines for each compound and outliers beyond ±1.5x the interquartile range defined by whiskers are shown but were not used for statistical comparisons. Letters indicate statistically significant differences among samples as determined by ANOVA followed by Tukey’s HSD test (P < 0.05).

**Figure 3.**
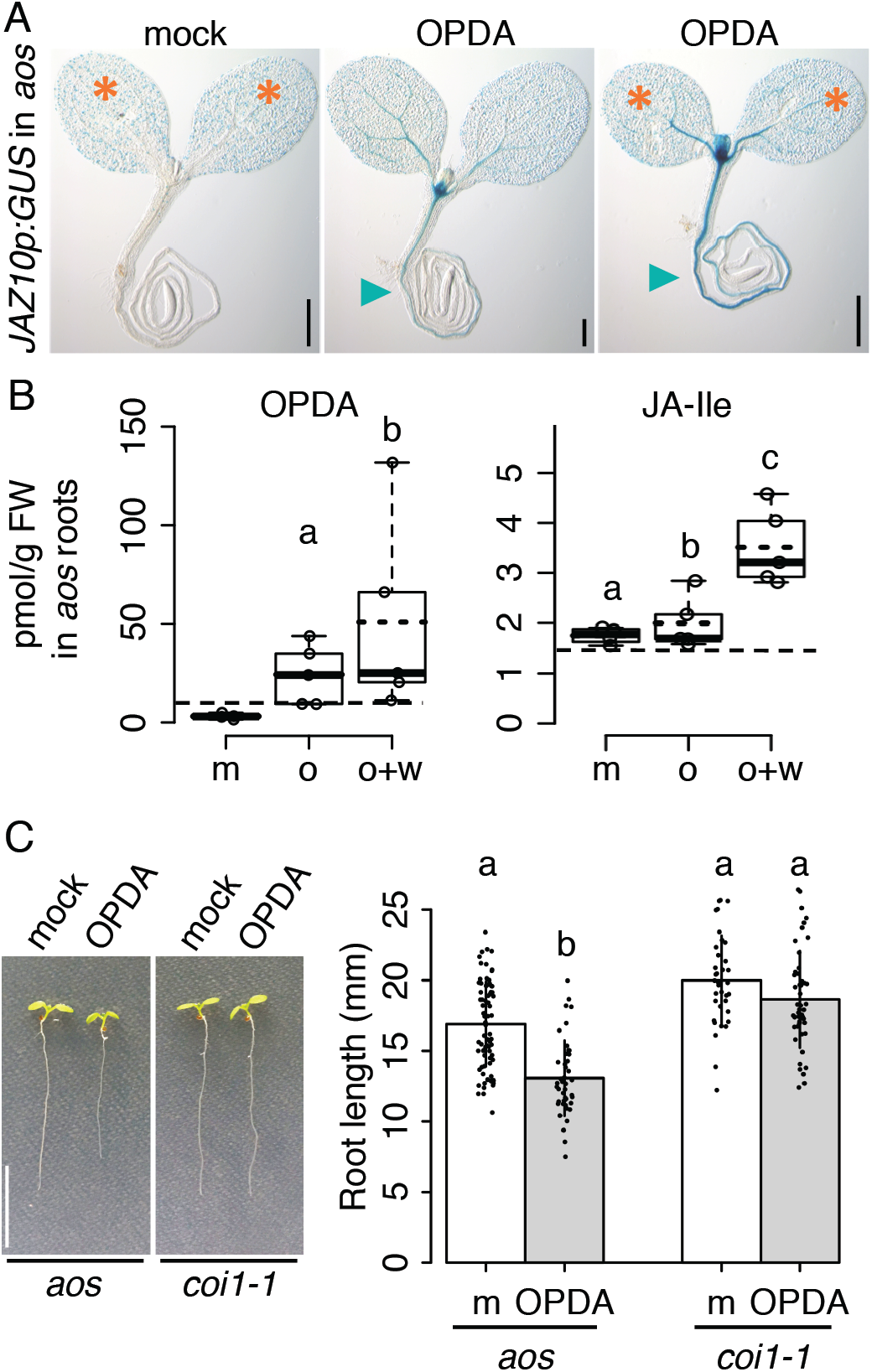
Exogenously applied OPDA can move basipetally from shoot to roots and inhibit root growth through COI1-mediated signaling. **(A)** Histochemical detection of *JAZ10p:GUS* activity in 5 d old *aos* seedlings 3 h after both cotyledons were treated with mock (cotyledon wounding + mock application); 30 μM OPDA; or 30 μM OPDA + wounding. OPDA was dissolved in agar to prevent leakage into roots. Wounding sites are indicated with orange asterisks and blue arrowheads depict reporter activation in roots. Each image is a representative sample of 18-20 biological replicates. Scale bars = 0.5 mm. **(B)** Box plot summary of OPDA and JA-Ile levels in roots of 14 d old *aos* plants 1 h after 5 leaves were wounded and treated with mock (m); treated with 30 μM OPDA (o); or wounded and treated with 30 μM OPDA (w+o). Circles depict biological replicates (4-5 per treatment), each consisting of roots from 15-20 individuals. Values below the limit of quantification (LOQ) indicated by dotted lines were not used for statistical comparisons. Note that all values for JA are below LOQ (Supplemental Figure 5). **(C)** Root length of 7 d old *aos* and *coi1-1* seedlings following exogenous application of mock (m) or 30uM OPDA in agar to above ground tissues for 48 h. Scale bar = 1cm. Bars show means of individual measurements represented by black dots (30-79 per treatment) ±SD. Letters indicate statistically significant differences among samples as determined by ANOVA followed by Tukey’s HSD test (P < 0.05).

To verify these findings, we followed *JAZ10* expression in mutants lacking phloem tissues in their primary roots and whose root vascular bundles are composed of sole protoxylem elements. We hence wounded cotyledons of the cytokinin (CK) histidine kinase CRE1/AHK4 receptor mutant *wooden leg* (*wol-1*) [32] and those of a line overexpressing the CK catabolic enzyme CK oxidase/dehydrogenase 7 (*35Sp:CKX7-GFP*) [33], to determine if their phloem-less primary roots were able to induce JA signaling in response to shoot injury. Although both mutants exhibit short roots due to compromised development of their root vascular system [32, 33], mechanical injury to their cotyledons induced *JAZ10* expression similar to wt levels (Fig. 1E). Contrarywise, while wt roots elicited *JAZ10* expression following shoot wounding, both CK mutants lacking phloem tissue in their primary roots almost entirely failed to induce JA signaling in this organ (Fig. 1E).

Taken together, our data indicate that a functional phloem is necessary and sufficient for wound-induced translocation of endogenous jasmonates into roots. The accumulation of JA and OPDA in vascular bundles and expression of JA biosynthesis enzymes in the phloem including companion cells, further support our findings on the phloem being the major tissue for oxylipin translocation to roots [34]. Interestingly, GLR-mediated leaf-to-leaf WASP signals require the concomitant participation of both phloem and xylem tissues to transmit damage signals and activate JA production in distal leaves [6, 7]. Hence, although hormone translocation and the rapid GLR-mediated long-distance signals likely involve different transmission mechanisms, their propagation routes overlap in the phloem.

### Endogenous OPDA is a mobile jasmonate species

To determine which oxylipin(s) move(s) basipetally from wounded shoots into undamaged roots through the phloem, we quantified root levels of JA-Ile and its precursors via targeted Ultra-Performance Liquid-Chromatography (UPLC)-electrospray ionization (ESI)-mass spectrometry (MS) [35]. In order to obtain sufficient root material required for the analysis and to verify if wound-induced shoot-to-root translocation of oxylipins occurs at later stages of plant development, we monitored *JAZ10p:GUS* expression in reciprocal grafts between the wt and the JA-deficient *aos* mutant in adult plants (Supplemental Figure 4). Wt scions grafted onto wt rootstocks (wt/wt) exhibited graft-induced basal reporter activity only near the graft junction which did not extend to above-or below-ground tissues. Leaf wounding caused strong reporter activation in both aerial tissues and rootstocks. Reporter expression was low in both control and wounded *aos*/*aos* grafts, and the *aos*/wt combination did not exhibit basal nor induced reporter activation at distal sites from the grafting site. Lack of reporter activation in rootstocks of scion-wounded *aos*/wt grafted plants indicated that 18:3/16:3 or 13-HPOT/11-HPHT generated in *aos* did not translocate to wt rootstocks to activate reporter expression, and further confirmed that WASP-mediated electrical signals are unlikely to regulate shoot-to-root induction of JA signaling. Conversely, although basal reporter activity was low in wt/*aos* plants, the *JAZ10p:GUS* reporter was strongly induced after leaf wounding in both wt scions and JA *aos* rootstocks (Supplemental Figure 4). Hence, wound-induced basipetal translocation of oxylipins observed in young seedlings [8] also occurs at later stages of plant development.

**Figure 4.**
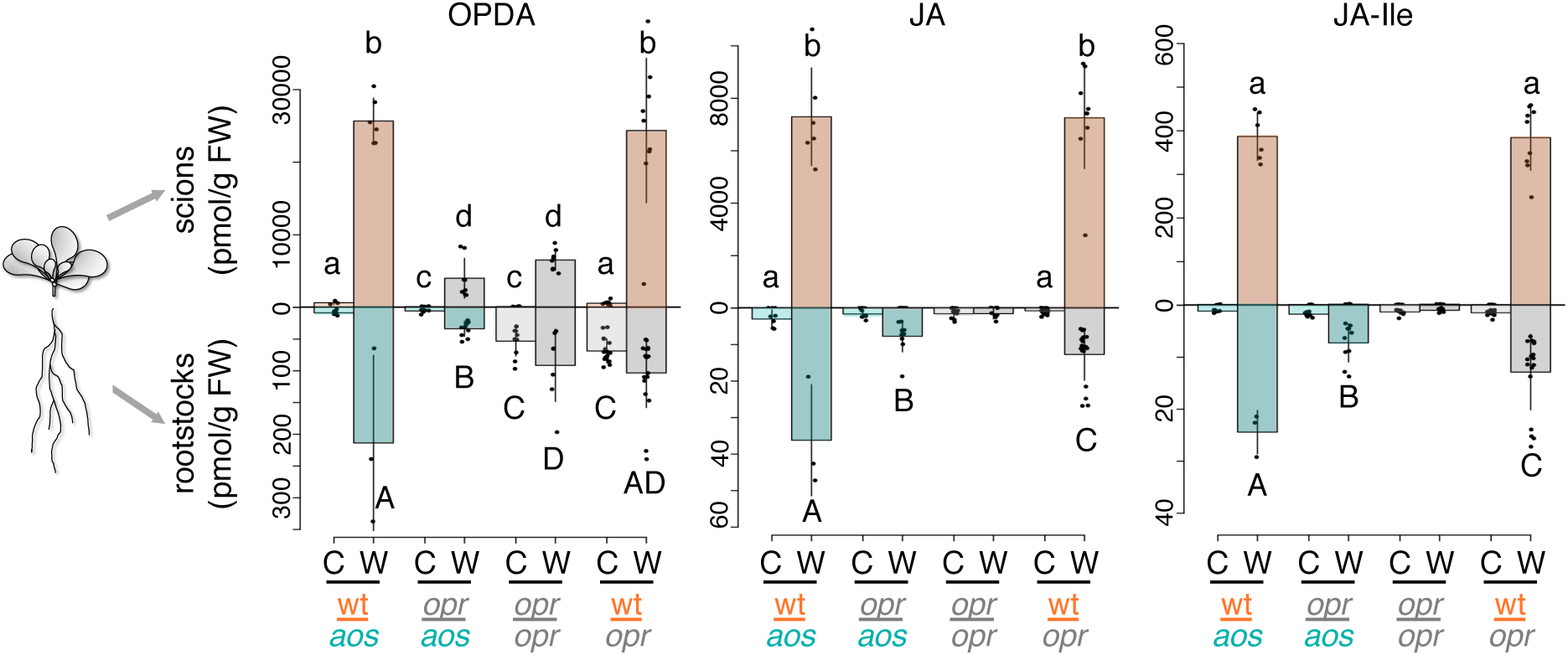
Shoot-derived OPDA is converted to JA and JA-Ile in undamaged roots. OPDA, JA, and JA-Ile levels in scions and rootstocks of grafted plants from indicated genotypes, where *opr* refers to the *opr2-1 opr3-3* double mutant deficient in all compounds downstream of OPDA. Quantification was performed at control conditions (C) or 1 h after shoot wounding (W). Bars show means of individual measurements represented by black dots (4-18 samples per treatment) ±SD, each consisting of pools from 3-5 scions or 15-20 rootstocks. Letters indicate statistically significant differences among samples as determined by ANOVA followed by Tukey’s HSD test (P < 0.05). Values below the LOQ were not used for statistical analysis.

As scion wounded wt/*aos* grafted plants still activated JA signaling in the JA-deficient rootstock, we reasoned that the rootstock would accumulate the translocating compound and its downstream derivatives. Due to their short half-life (<20 s at 0 C°) and high instability, the plastidial OPDA precursors allene oxides 12,13-EOT and 10,11-EHT were not considered as possible translocating compounds [11] (Supplemental Figure 1). Analyzed compounds included the most abundant JA species produced after wounding (OPDA, JA and JA-Ile) [9], whereas β-oxidation intermediates (OPC-8, OPC-6 and OPC-4) were below the limit of quantification (LOQ) in our system. We first quantified root oxylipin levels in ungrafted wt and *aos* plants under basal conditions and 1 h after shoot wounding (Fig. 2). As expected [36], *aos* roots did not accumulate any basal or wound-induced oxylipin, and wt roots exhibited higher basal OPDA levels (36.5±18 pmol/g FW) compared to downstream compounds, which increased further after wounding (68.3±24.8 pmol/g FW). JA and JA-Ile levels were very low in wt unwounded roots and increased significantly upon shoot wounding.

Next, OPDA and JA levels in rootstocks of control and shoot-wounded wt/wt grafted plants were similar to those of ungrafted wt plants, whereas wound-induced JA-Ile levels were lower (Fig. 2). This suggested that the graft junction might negatively impact bioactive hormone accumulation. Rootstock profiling from *aos*/wt combinations revealed that oxylipin levels do not increase in wt rootstocks in response to damage of *aos* scions, strengthening previous findings that root activation of JA biosynthesis following shoot injury is entirely dependent on hormone translocation [8]. Finally, OPDA, JA and JA-Ile levels were very low in unwounded wt/*aos* rootstocks but, all three compounds increased significantly upon wt scion wounding. Due to the high instability of the allene oxide OPDA precursors [11], we concluded that OPDA measured in *aos* rootstocks translocated from wounded wt scions. Moreover, basal OPDA levels in rootstocks of wt/*aos* grafts were undetectable as in ungrafted *aos* rootstocks, indicating that the shoot-to-root translocation of endogenous OPDA occurred only after scion wounding and not constitutively (Fig. 2). In addition to OPDA, the data showed increased levels of JA and JA-Ile in the *aos* rootstocks, suggesting that both compounds may also translocate from wounded wt scions or derive from OPDA conversion directly in the *aos* rootstock. Nonetheless, JA amounts were lower in wounded wt/*aos* rootstocks compared to wt or wt/wt controls. Previous reports analyzing endogenous oxylipin translocation excluded leaf-to-leaf JA and JA-Ile movement [3], or concluded that JA but not JA-Ile is moving acropetally to undamaged tissues [26, 27]. However, OPDA was not analyzed in those experiments, and after re-evaluation of the published material, the data is also compatible with OPDA translocation.

### Exogenously applied OPDA can translocate from wounded shoots into undamaged roots

Exogenous OPDA is able to activate gene expression by its conversion to the bioactive hormone [14, 37]. To support our findings on endogenous OPDA mobility, we applied exogenous OPDA to above-ground tissues of the OPDA-deficient *aos* mutant and monitored the consequences in roots via two separate assays. First, *JAZ10p:GUS* reporter analysis showed that compared to mock controls, OPDA treatment induced a slight increase in JA signaling predominantly in the vasculature at the sites of application as well as in below-ground tissues (Fig. 3A). Overall reporter activation was considerably stronger when OPDA application was coupled to cotyledon wounding, suggesting that wounding facilitates OPDA penetration and/or tissue damage stimulates shoot-to-root OPDA translocation. Next, we profiled oxylipins in *aos* roots following OPDA applications to shoots (Fig. 3B). Although JA levels were below the LOQ in all measurements (Supplemental Figure 5), OPDA and JA-Ile increased slightly after OPDA application compared to the mock, and they increased further when OPDA treatment was combined with cotyledon wounding. Hence, both endogenous and exogenously applied OPDA can translocate to roots following shoot wounding to activate JA signaling.

**Figure 5.**
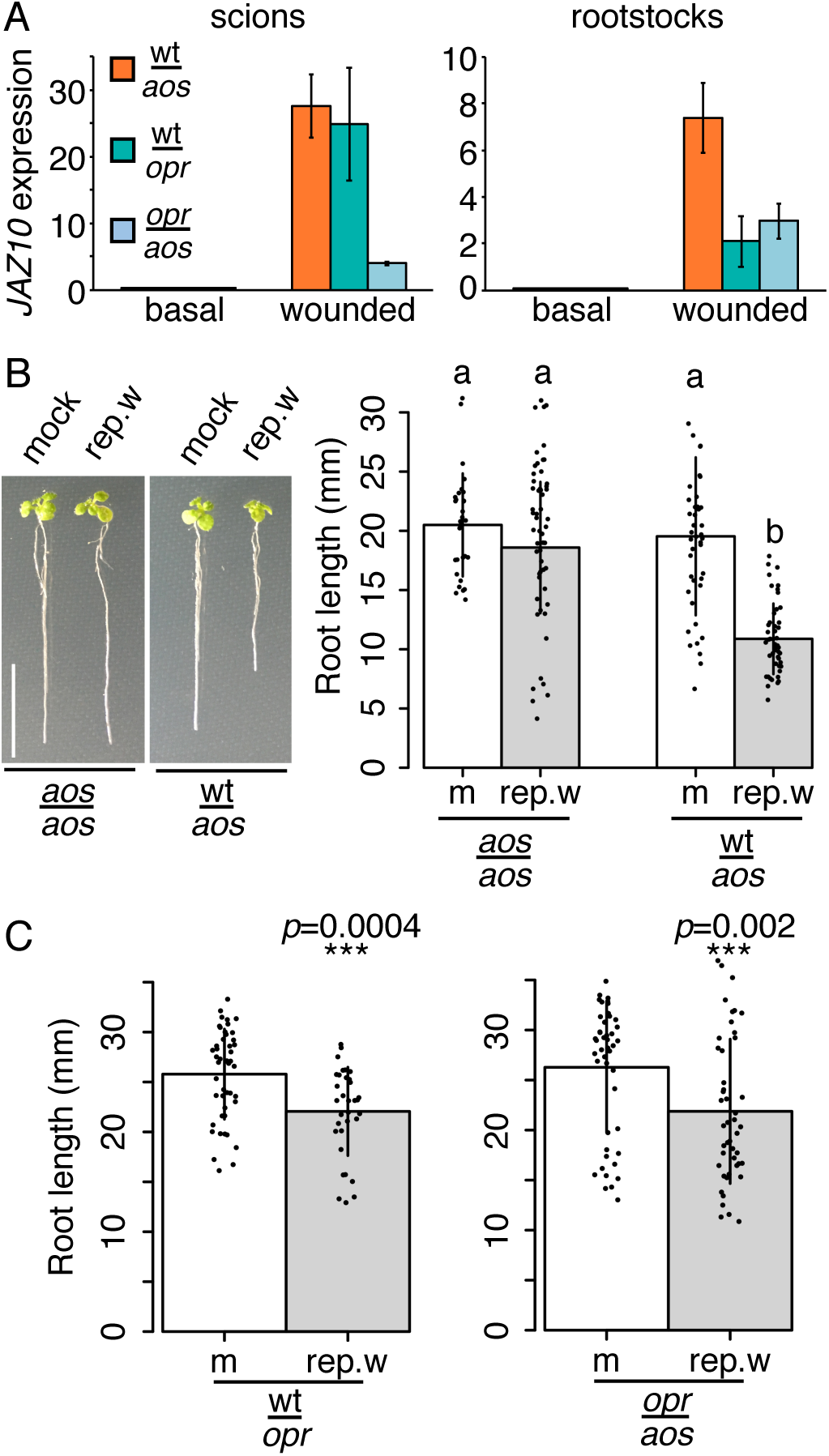
Shoot-to-root translocation of OPDA is essential to induce full JA root signaling and coordinate whole plant growth responses. **(A)** qRT-PCR of *JAZ10* expression in scions and rootstocks of the indicated genotypes, where *opr* refers to *opr2-1 opr3-3*, under basal conditions and 1 h after extensive shoot wounding. *JAZ10* transcript levels were normalized to those of *UBC21*. Bars represent the means of three biological replicates (±SD), each containing a pool of organs from five 3-week-old grafted plants. **(B-C)** 12 d old grafted plants of indicated genotypes grown under mock (m) conditions or subjected to repetitive shoot wounding (rep.w), and relative bar charts showing means of individual measurements represented by black dots (30-79 per treatment) ±SD. Scale bar = 1cm. Letters in **B** denote statistically significant differences among samples as determined by ANOVA followed by Tukey’s HSD test (P < 0.05), and stars in **C** depict significant Student’s *t*-test results with relative p-values (p).

To determine if the translocation of OPDA requires conversion into the bioactive hormone to induce JA responses, we treated cotyledons of *aos* seedlings with exogenous OPDA and measured its effects on primary root length (Fig. 3C). Two days after application, roots of OPDA-treated *aos* seedlings were significantly shorter (13.1 ± 2.8 mm) than mock controls (16.9 ± 3 mm), indicating that shoot-derived OPDA could inhibit root growth. Importantly, *coi1-1* seedlings were insensitive to the treatment, implying that OPDA must be converted to the bioactive hormone to exert its physiological function. Although several studies proposed COI1-independent signaling roles for OPDA, eg. [37-39], our data suggests that shoot-derived OPDA induces JA signaling and represses root growth via conversion into JA-Ile and subsequent COI1-signaling.

### Translocated OPDA is converted to JA and JA-Ile in undamaged roots

Recently, a double knockout mutant in *OPR3* and *OPR2* was found to entirely abolish JA and JA-Ile production [14], while retaining the capacity to synthesize OPDA (Supplemental Figure 1). We thus employed the *opr2-1 opr3-3* double mutant (hereafter abbreviated to *opr*) to quantify the amount of shoot-derived OPDA translocating into roots and its subsequent conversion to JA and JA-Ile in this organ. To account for the positive feedback loop where the activation of JA signaling stimulates its own biosynthesis [40, 41] present in the wt but absent in *opr*, the experimental setup included hormone profiles from scions. As expected (Fig. 2) [42], basal levels of OPDA, JA and JA-Ile were low in both scions and rootstocks of wt/*aos* control plants and they increased significantly in both organs following scion wounding (Fig. 4). Both basal and wound-induced OPDA scion levels were lower than wt in the *opr*/*opr* control grafts and, importantly, they increased significantly in both organs following shoot wounding. Wound-triggered OPDA increase in *opr*/*opr* plants did not produce measurable JA or JA-Ile levels, however wounding of *opr*/*aos* combinations resulted in the sole increase of OPDA in *opr* scions and the induction of all compounds (OPDA, JA, JA-Ile) in *aos* rootstocks (Fig. 4). These findings demonstrated that shoot-derived OPDA is converted to JA and JA-Ile directly in the undamaged root. In spite of ~5-fold reduced OPDA levels available for long distance translocation in wounded *opr* versus wt scions (4 ± 2.7 vs. 25.7 ± 3 nmol/g FW respectively), a proportionate OPDA amount was detected in *aos* rootstocks (33.6 ± 12 pmol/g FW from *opr* vs. 214 ± 138 nmol/g FW from wt scions) and the resulting conversion into bioactive JA-Ile followed a similar trend (7.2 ± 3.7 pmol/g FW from *opr* vs. 24.4 ± 4.2 nmol/g FW from wt scions). Hence, it is likely that during the wound response of wt plants, OPDA translocating to roots generates higher levels of JA-Ile than those measured in *opr*/*aos*.

In addition to OPDA, scion wounding of wt/*aos* grafts resulted in increased JA and JA-Ile levels in *aos* rootstocks (Fig. 2). To verify if OPDA is the sole JA species translocating into roots after shoot damage or if other compounds downstream of OPDA are also mobile, we quantified oxylipins in wt/*opr* grafted plants (Fig. 4). As *opr* is unable to synthesize de novo JA and JA-Ile, the increase of those compounds in *opr* rootstocks after wt scion damage indicated that additional compounds downstream of OPDA translocate basipetally from wounded shoots. OPC-8, OPC-6, OPC-4, JA and JA-Ile are thus all possible transported candidates, and endogenous JA and JA-Ile were found to be mobile in tomato and wild tobacco [26, 27]. The identification of NPF2.10/GTR1 as JA and JA-Ile transporter [43, 44] and of AtABCG16/JAT1 mediating nuclear JA-Ile influx [45], further underline the importance of a regulated distribution of jasmonate species across cells and organs. Due to unidentified multiple enzymes [15] and gene redundancy (Supplemental Figure 1), the unavailability of genetic stocks depleted in OPDA derivatives at each step of JA-Ile biosynthesis downstream of *opr2 opr3* prevented us from dissecting the mobility of OPDA derivative(s) further. Nevertheless, our data indicated that OPDA translocation accounts for a large proportion (at least 30%) of JA-Ile production in unwounded roots, as rootstock JA-Ile levels were 24.4 ± 4.2 pmol/g FW in WT/*aos*, 7.2 ± 3.7 pmol/g FW in *opr*/*aos* and 12.8 ± 7 pmol/g FW in WT/*opr* following wounding of the relative scions.

### OPDA translocation from wounded shoots is essential to induce full JA root signaling and coordinate whole plant growth responses

Given that both OPDA and its derivative(s) are mobile compounds, we next assessed what are their contributions in activating root JA signaling and in regulating JA-mediated growth responses following shoot damage. To this end, we used wt/*aos* as control grafts to determine the effects of both OPDA and its derivative(s), wt/*opr* combinations for OPDA derivative(s), and *opr*/*aos* for evaluating the effects of OPDA only. Transcript levels of the JA marker *JAZ10* were low in both scions and rootstocks of wt/*aos* grafted plants and increased significantly in both organs upon scion wounding (Fig. 5A). Wound-induced *JAZ10* transcripts reached similar levels in scions of wt/*opr* grafts, but were two thirds lower than those of wt/*aos* grafts indicating that the translocation of OPDA derivative(s) into undamaged roots account for approximately 30% of JA signaling increase. Remarkably, although scion wounding triggered no JA-Ile accumulation and very little *JAZ10* induction in *opr*/*aos* scions, shoot derived OPDA was sufficient to trigger JA signaling in *aos* rootstocks and induce *JAZ10* transcript levels to 40% of those found in wt/*aos* grafts (Fig. 4-5).

To uncover if in addition to inducing transcriptional changes (Fig. 5A, Supplemental Figure 4), the translocating compounds have tangible effects on root physiology, we quantified primary root lengths from grafted plants subjected to repetitive shoot wounding, a treatment that is known to stunt root growth in a JA-dependent manner [36]. First, we verified if the method was applicable to grafted plants, which may exhibit variations in their primary root length caused by grafting and transfer (Fig. 5B). In spite of the data variability, root lengths of scion wounded *aos*/*aos* grafts (18.6 ± 5.5 mm) were not significantly different from the respective mock controls (20.5 ± 4.3 mm). Conversely, scion wounded wt/*aos* plants exhibited significantly shorter roots (10.9 ± 3 mm) compared to the unwounded mocks (19.5 ± 6.7 mm). Thus, grafted plants are amenable to repetitive shoot wounding to evaluate the JA-dependent inhibition of root growth. Both translocated OPDA and translocated OPDA derivative(s) participated in repressing root growth after shoot damage, as rootstocks from scion wounded wt/*opr* and *opr*/*aos* plants exhibited similar reductions in their primary root lengths (Fig. 5C). Taken together, our data indicate that the translocation of OPDA and its downstream compound(s) are essential and cannot be compensated by one another to induce full JA root signaling and coordinate shoot-to-root growth responses. Nonetheless, as several OPDA derivatives may be mobile accounting collectively for the observed phenotypes (eg. Fig. 5), and because *opr* scions generate lower OPDA amounts available for shoot-to-root translocation, the data in fact suggest that OPDA might be a major form of translocating JA species coordinating wt shoot-to-root responses.

Overall, our work shows that the JA-Ile precursor OPDA is an important communication molecule that enables plants to respond to shoot derived stimuli and coordinate plant architecture. Probably due to its high lipophilicity, previous studies aimed at identifying mobile jasmonates did not systematically investigate OPDA as a possible mobile compound [3, 21, 22, 24-27]. In fact, OPDA has a higher octanol-water partition coefficient compared to JA and JA-Ile [46], and thus it remains unknown how can this molecule travel long-distances to eventually activate JA signaling. Notably, OPDA, but not JA nor JA-Ile, is present in early land plants such as bryophytes and its levels increase after wounding [46]. It is possible that the emergence and development of an OPDA transport system had a longer evolutionary time to attain efficiency. Findings presented here open up new perspectives regarding the mechanisms of OPDA translocation, including the search for cellular OPDA exporters, putative carriers within the vascular stream and OPDA importers in distantly located cells. Furthermore, our results highlight the importance of a wound-regulated distribution of endogenous oxylipins to effectively accomplish bioactive hormone functions, and pave the way to identify the molecular mechanisms of their transport in vascular plants.

## METHODS

### Plant material and growth conditions

Previously described *Arabidopsis thaliana* wild-type (wt) and JA-deficient *aos* lines [10], both in the Columbia (Col) background harboring the *JAZ10p:GUS* reporter [8]; *coi1-1* [47]; *wol-1* [32] from NASC; *35Sp:CKX7* [33]; and *opr2-1 opr3-3* [14] were used throughout this study. After seed sterilization and stratification for 2 d at 4°C, plants were grown aseptically on 0.5X solid Murashige and Skoog medium (MS; 2.15 g/L, pH 5.7; Duchefa) supplemented with 0.5 g/L MES hydrate (Sigma) and 0.7% or 0.85% (w/v) agar (Plant Agar for cell culture, AppliChem GmbH), for horizontally or vertically grown plants respectively. Horizontally grown seedlings were germinated on a nylon mesh placed on top of the MS media as described [30]. Controlled growth conditions were set at 21°C under 100 μE m^-2^ s^-1^ light, with a 14-h-light/10-h-dark photoperiod.

### Mechanical wounding

Cotyledon wounding of seedlings was performed as described [30]. For single and triple primary root wounding, roots of 5 d old vertically grown seedlings were gently squeezed with fine (4.5 S) forceps under a stereomicroscope in the middle of the root (single) or at approximatively three equidistant portions along the root (triple). For oxylipin profiling, qRT-PCR from grafted plants and histochemical detection of GUS activity in vertically grown adult plants, rosettes were wounded when their roots reached the bottom of the vertical plate (3-week-old plants). Each plant was wounded with serrated 18/8 forceps by squeezing the surface of all (6-7) leaves extensively. Repetitive scion wounding in grafted plants was initiated immediately after graft evaluation (8 d after grafting) and transfer to vertical MS media plate. Plants were pierced with a 36-gauge beveled NanoFil needle (110 μm outer diameter) on one leaf in aseptic conditions under a stereomicroscope as described [36]. Wounding started in the morning (7– 8 am) and was repeated every 12 h, for a total of 4 d and 8 wounds per plant. Primary root length measurements were done as described [30]. In all cases, wounded and mock-treated plants were returned to controlled conditions for the indicated times prior harvesting.

### Micrografting

Grafting was performed as described [31]. Graft formation was evaluated at 8 d by the attachment of scion to rootstock without the development of adventitious roots. Successful grafts were transferred to vertical MS medium (5 plants per vertical plate) and grown further depending on the application.

### Phloem and xylem connectivity assays

Assays were performed 2-8 d after grafting as described [31]. For phloem reconnection assays, plants were treated in the grafting dish. Cotyledons were squeezed with sharp forceps and 1 μl of 1 mM 5(6)-Carboxyfluorescein diacetate (CFDA) dissolved in melted (60°C) 0.8% agar was immediately pipetted onto the cotyledon where it solidified. For xylem reconnection assays, grafts were cut below the root-hypocotyl junction, and hypocotyls were then placed in solidified 1 mM CFDA-agar without scions touching the agar. After 1 h (phloem assays) or 20 min (xylem assays) incubation, the fluorescence signal in the rootstock vasculature (phloem assay) or scion (xylem assay) was evaluated with a Nikon AZ100 fluorescence microscope fitted with a GFP filter and imaged with a Nikon Digital Sight DS-5Mc camera. For phloem reconnection assays and histochemical detection of GUS activity from the same sample, grafted plants were wounded with a 25G x 5/8’’ needle (0.5 mm x 16 mm) 2 h prior CFDA treatment. After CFDA imaging in the rootstock, samples were assayed for GUS activity.

### Histochemical detection of GUS activity

Plants were wounded 3 h before prefixing with 90% acetone for 1 h. Samples were rinsed with 50 mM NaHPO_4_ buffer (pH 7), and subsequently stained with staining solution (10 mM EDTA pH 8.5, 50 mM NaHPO_4_ pH 7.0, 3 mM K_3_Fe(CN)_6_, 3 mM K_4_Fe(CN)_6_, 0.1% Triton X-100, 0.5 mg/mL X-Gluc). Samples were vacuum infiltrated and then incubated at 37°C in the dark for either 4 h (seedlings) or overnight (adult plants). The reaction was stopped by replacing the solution with 70% EtOH. For seedling imaging, samples were mounted in clearing solution (8 g chloral hydrate: 2 ml glycerol: 1 ml water) and photographed with a Leica M165 FC stereomicroscope fitted with a Leica MC170 HD camera. Adult grafted plants were imaged on a Nikon SMZ1270 stereomicroscope fitted with a Nikon Digital Sight DS-Fi2 camera and bottom illumination from a Transotype screen while kept in 70% EtOH.

### RNA isolation and qRT-PCR assays

RNA extraction and gene expression of *JAZ10* (At5g13220) and *UBC21* (At5g25760) was performed as described [36].

### Exogenous OPDA applications

Seedlings (5 d old) or adult plants (vertically grown, 3-week-old) were wounded on both cotyledons (seedlings) or on 5 rosette leaves (adult plants) with a 25G x 5/8’’ needle (0.5 mm x 16 mm) under a dissecting microscope immediately before placing 2 μL of mock or 2 μl of OPDA (30 μM in 0.8% 60°C melted agar) solution on each wounding site (tot of 4 μL per seedling or 10 μL per adult plant). Shoot applications were done in agar in order to prevent cross-contamination and leakage of solutions to roots. Unwounded plants were treated with the OPDA solution only. OPDA (*cis*-12-oxo-phytodienoic acid) was synthesized enzymatically according to [48]. Samples were incubated under controlled growth conditions for 3 h (histochemical GUS detection in seedlings) or 1 h (oxylipin quantification in roots by LC-MS) before harvest. For measuring primary root lengths, mock or 30 μM OPDA treatment (as above) was applied on shoots of 5 d old vertically-grown seedlings immediately after wounding both cotyledons to increase treatment penetration, and root lengths were measured 48 h later with the ImageJ software (http://rsb.info.nih.gov/ij/).

### OPDA, JA and JA-Ile quantification

Control and wounded adult plants were harvested just before roots reached the bottom of the vertical plates (12-16 d after grafting or 15-18 d for un-grafted plants). Roots were excised beneath the hypocotyl/root junction and hypocotyls were discarded from shoots immediately prior freezing in liquid N_2_ (flash freezing) [42]. The time from cutting to flash freezing was monitored and kept below 16 s. Phytohormone measurements were performed on a minimum of 4 biological replicates, each consisting of pooled roots from 15-20 plants and pooled shoots from 3-5 plants, yielding approximately 50 mg of fresh weight. Extraction and quantitative analysis of oxylipins was performed as described [35], with minor modifications. Deuterium labeled internal standard [^2^H_5_]*cis*-12-oxo-phytodienoic acid ([^2^H_5_]OPDA) was prepared from (17-^2^H_2_,18-^2^H_3_)-linolenic acid as in [48]; [^2^H_6_] jasmonic acid ([^2^H_6_]JA) was obtained as described in [49]; and [^2^H_2_]N-(-)-jasmonoyl isoleucine ([^2^H_2_]JA-Ile) was prepared from (^2^H_2_)-JA and Ile according to [50]. Frozen plant material was homogenized with 250 μL of methanol containing 0.5 ng of [^2^H_6_]JA and [^2^H_2_]JA-Ile, and 1 ng of [^2^H_5_]OPDA in a Tissue Lyzer. After centrifugation at 10,000 rpm for 7 min, the supernatant was transferred to 2 ml tubes (Sarstedt) containing 1.7 ml milliQ H_2_O to polarize the solution. Samples were then subjected to solid-phase extraction (SPE) in 96-well plates cation exchange columns (Macherey-Nagel HR-XC), pre-conditioned with 500 μl methanol and equilibrated with 500 μl milliQ H_2_O. All centrifugation steps were carried out at 500 rpm for 5-7 min. Oxylipins were eluted with 900 μl acetonitrile and concentrated to 100 μl via vacuum evaporation before injection into the Ultra-Performance Liquid-Chromatography (UPLC)-electrospray ionization (ESI)-mass spectrometry (MS) system. Separations and analysis were performed as described [35], but on a QTrap 6500 system. The limit of quantification (LOQ = 3x limit of detection) was determined from an Arabidopsis matrix as 7 pmol/g FW for OPDA, 13.2 pmol/g FW for JA and 1.5 pmol/g FW for JA-Ile. Measurements below these values were not considered for statistical analyses.

### Statistical analysis

Box plots, multiple comparisons [analysis of variance (ANOVA) followed by Tukey’s honest significant difference (HSD) test] and correlation coefficient analysis for association between paired samples followed by Person product moment correlation coefficient p-value were performed in R 3.4.4.

## SUPPLEMENTAL DATA

**Supplemental Figure 1.** JA-Ile biosynthesis in Arabidopsis

**Supplemental Figure 2.** Phloem and xylem reconnection assays and *JAZ10p:GUS* reporter analysis in wt/*aos* grafts.

**Supplemental Figure 3.** The GUS protein is not translocating across the graft junction following scion wounding.

**Supplemental Figure 4.** Basipetal shoot-to-root transport of JA species in adult Arabidopsis plants.

**Supplemental Figure 5.** Box plot summary of JA levels in *aos* roots following exogenous OPDA application to shoots.

## ACKNOWLEDGEMENTS

We thank R. Solano and A. Chini for *opr2-1 opr3-3* seeds; T. Schmuelling and I. Koellmer for *35Sp:CKX7* seeds; O. Miersch for providing OPDA and deuterated standards; and R. Dreos for R expertise.

