## Supplemental Figure for "Shoot-to-root translocation of the jasmonate precursor 12-oxo-phytodienoic acid (OPDA) coordinates plant growth responses following tissue damage"

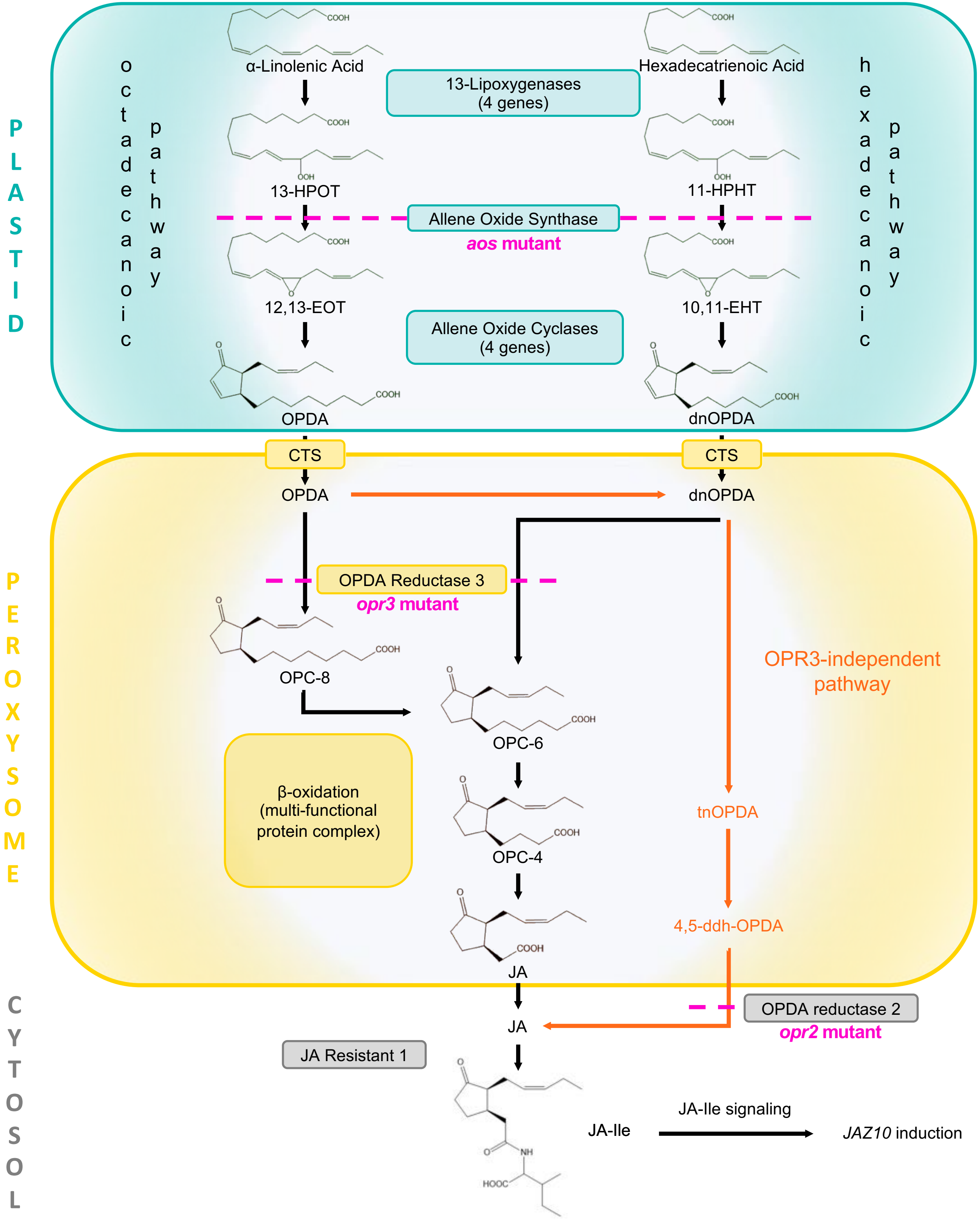


**Supplemental Figure 1. JA-Ile biosynthesis in Arabidopsis** starts from de-esterification of plastidial membrane lipids (mainly galactolipids) to yield linolenic acid (18:3) and hexadecatrienoic acid (16:3). The OPDA, JA and JA-Ile deficient mutant *aos* is shown in red, and the JA and JA-Ile deficient *opr2 opr3* double mutant is indicated in magenta. Abbreviations: 13-HPOT (13(S)-hydroperoxy-octadecatrienoic acid), 11-HPHT (11(S)-hydroperoxy-hexadecatrienoic acid), 12,13-EOT ((13S)-12,13-epoxy-octadecatrienoic acid), 10,11-EOT ((11S)-10,11-epoxy-octadecatrienoic acid), OPDA (12-oxo-phytodienoic acid), dnOPDA (dinor-oxo-phytodienoic acid), CTS (ABC-type transporter COMATOSE), OPC-8 (3-oxo-2-(2-(Z)-pentenyl)-cyclopentane-1-octanoic acid), OPC-6 (3-oxo-2-(2-(Z)-pentenyl)-cyclopentane-1-hexanoic acid), OPC-4 (3-oxo-2-(2-(Z)-pentenyl) cyclopentane-1 butanoic acid), OPR (OPDA Reductase 3), tnOPDA (tetranor-OPDA), 4,5-ddh-OPDA (4,5-didehydro JA), JA ((+)-7-iso-jasmonic acid), JA-Ile ((+)-7-iso-jasmonoyl-isoleucine).

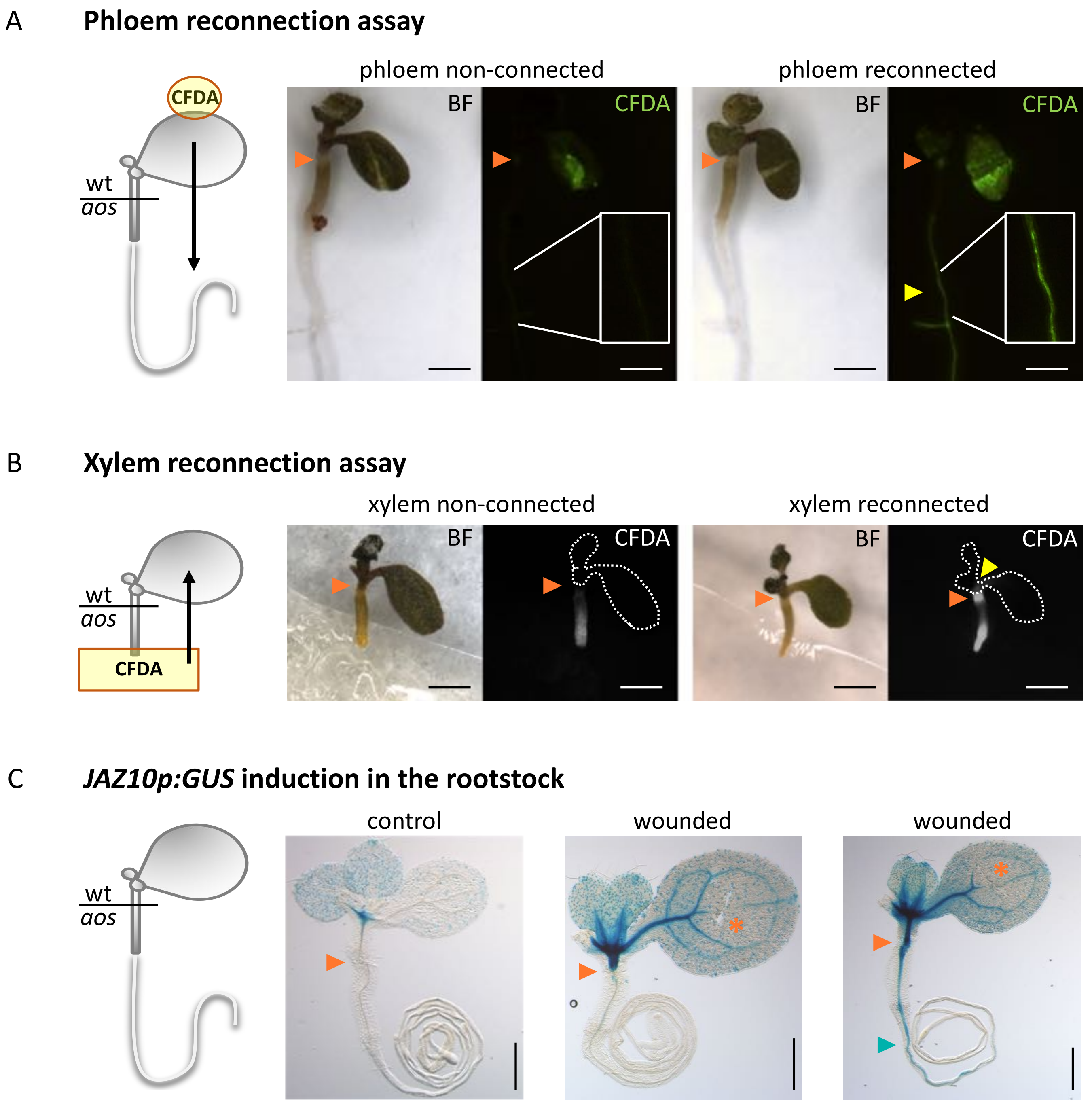


**Supplemental Figure 2. Phloem and xylem re-connection assays and *JAZ10p:GUS* reporter analysis in wt/*aos* grafts.** Seedling at 5 d post grafting are shown as examples. **a**, Following grafting, phloem connectivity is evaluated with carboxyfluorescein diacetate (CFDA) application on the scion and visualization of fluorescence in the rootstock. In the grafting dish, cotyledons are gently squeezed with fine forceps under a stereomicroscope and 1 µL of 1 mM CFDA in melted agar is immediately applied to the wound site. Absence or presence of fluorescence in the rootstock is evaluated 1 h after CFDA application as an indication of no phloem re-connection or successful phloem re-connection respectively. **b**, Xylem re-connection after grafting is evaluated by placing seedlings cut at the hypocotyl/root junction in solidified agar containing 1 mM CFDA. Absence or presence of fluorescence in the scion is monitored as an indication of xylem connectivity. **c**, *JAZ10p:GUS* reporter evaluation in *aos* rootstocks following wt scion wounding. BF refers to the bright field image, orange arrowheads indicate graft junctions, yellow arrowheads indicate CFDA florescence in the (**a**) rootstock or (**b**) scion, orange asterisks indicate wounding sites and blue arrowheads indicate *JAZ10p:GUS* reporter activity in the rootstock. Scale bars = 1 mm.

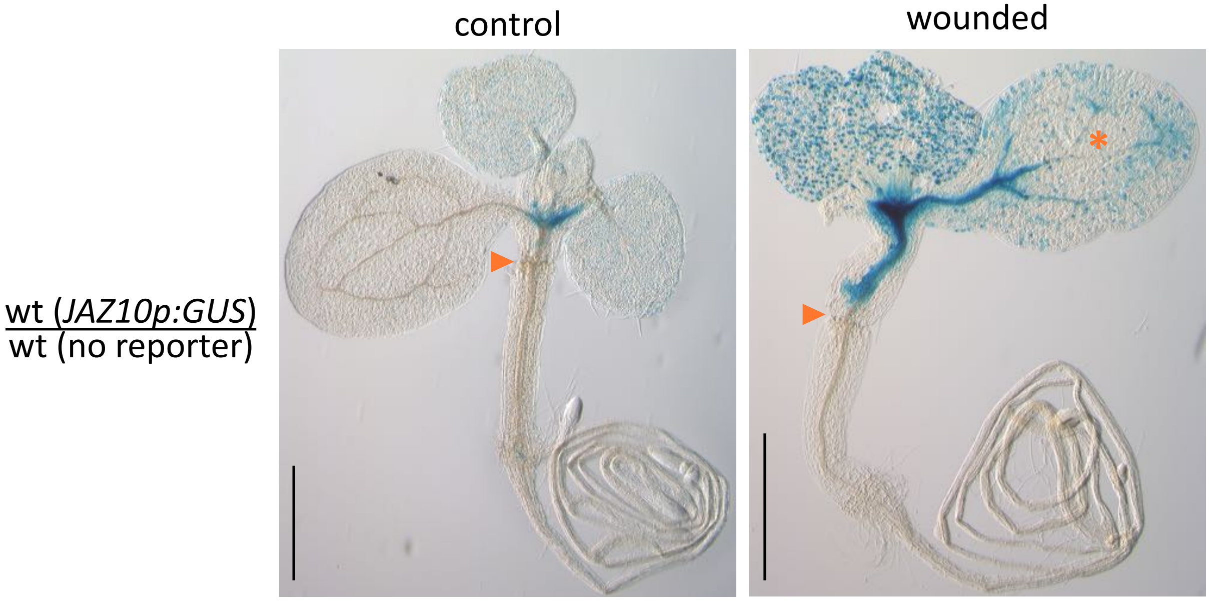


**Supplemental Figure 3. The GUS protein is not translocating across the graft junction following scion wounding.** Histochemical detection of GUS activity in 13 d old grafted seedlings of wt scions expressing the *JAZ10p:GUS* reporter onto wt (Col) rootstock without the reporter. GUS staining was performed 3 h after wounding 1 cotyledon as indicated by the orange asterisk. Note the lack of GUS activity in the rootstock of the wounded plant. Orange arrowheads indicate graft junctions. Each image is a representative sample from 18-20 biological replicates. Scale bars = 1 mm.

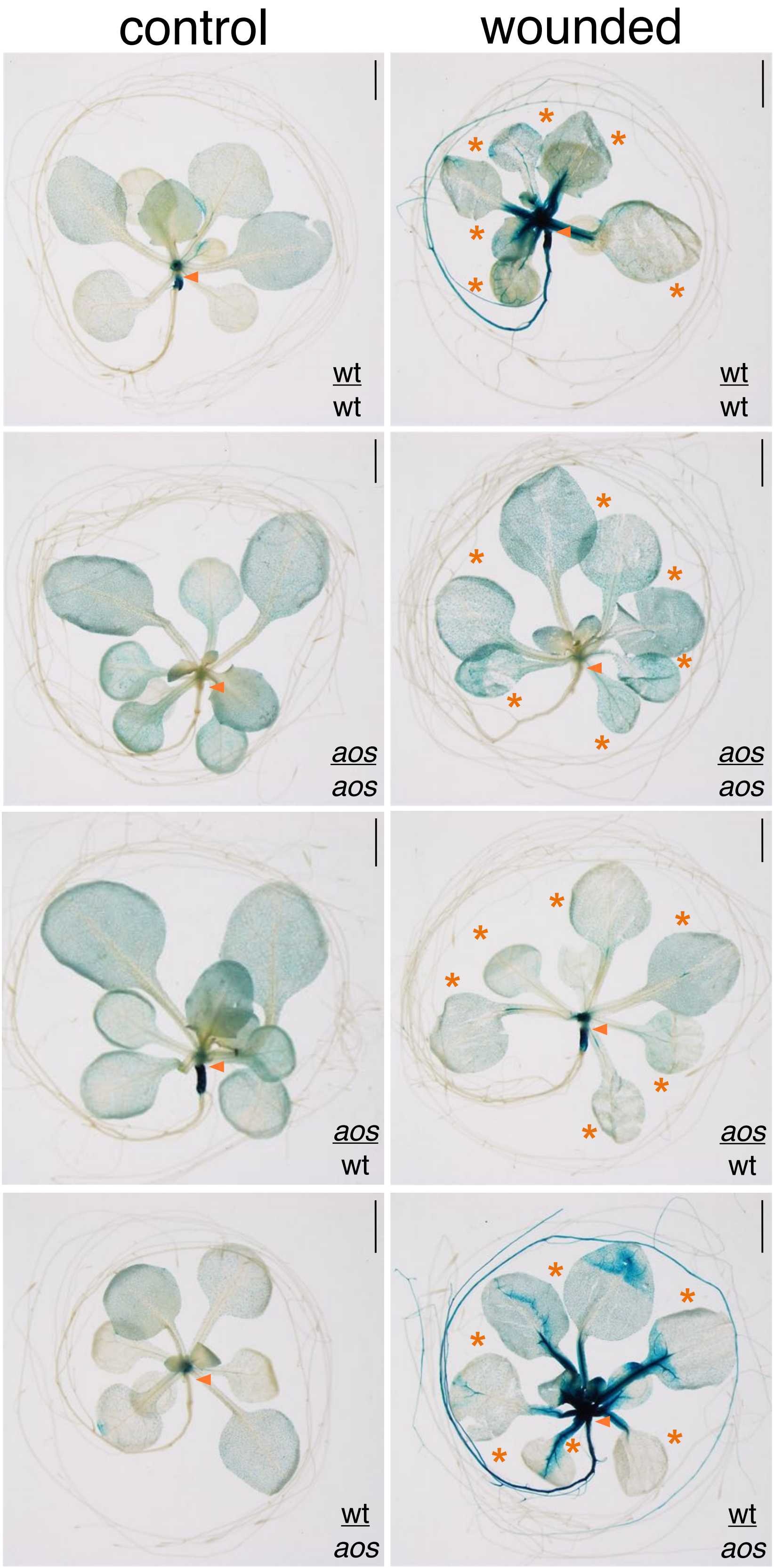


**Supplemental Figure 4. Basipetal shoot-to-root transport of JA species is occurring in adult *Arabidopsis* plants.** Histochemical detection of *JAZ10p:GUS* reporter activity in 3-week-old grafted plants of the indicated scion/rootstock genotypes: wt/wt, *aos*/*aos*, *aos*/wt, and wt/*aos* grafts. GUS staining was performed 3 h after extensively wounding adult leaves with serrated forceps at sites indicated by orange asterisks. Orange arrowheads indicate graft junctions. Each image is a representative sample from 8-11 biological replicates. Scale bars = 2 mm.

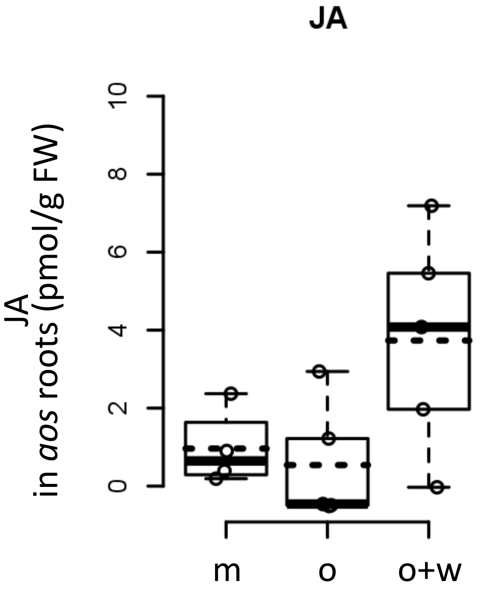


**Supplemental Figure 5.** Box plot summary of JA levels in roots of 14 d old *aos* plants 1 h after 5 leaves were wounded and treated with mock (m); treated with 30 µM OPDA (o); or wounded and treated with 30 µM OPDA (w+o). Circles depict biological replicates (4-5 per treatment), each consisting of roots from 15-20 individuals. Note that all values are below LOQ.
